# COGNITIVE DETERIORATION IS REVERSED BY AN INSULIN-LIKE GROWTH FACTOR 1 SENSITIZER IN A MOUSE MODEL OF ALZHEIMER DISEASE

**DOI:** 10.64898/2026.08.01.742197

**Authors:** J.A. Zegarra-Valdivia, M.Z. Khan, M. Vega, I. Torres Aleman

## Abstract

Previous observations in preclinical and clinical studies indicate a beneficial effect of insulin-like growth factor 1 (IGF-1) in different neurological illnesses, including Alzheimer’s disease (AD). AD is the most important neurodegenerative disease in the world and despite previous intensive research and the recent approval of putative disease-modifying therapies, available treatments provide only modest clinical benefit and do not halt disease progression. Consequently, there remains a pressing need to develop novel therapeutic strategies for AD. Since resistance to IGF-1 may be involved in development of AD, as it regulates cognition and amyloid β (Aβ) metabolism, we recently developed a small molecule IGF-1 sensitizer, AIK3a305, that crosses the blood brain barrier (BBB) and exerts modulatory actions in the brain. Using a mouse model of familial AD, the APP/PS1 mouse, we administered them AIK3a305 for 3 months. Treatment started at 12 months of age, when the disease is already well established, and cognitive deterioration readily measurable. One month after starting daily intraperitoneal injections of AIK3a305, mice showed normal cognitive performance in the Y maze, a measure of working memory that enables daily life activities. After 3 months, cognition remained fully preserved, mood-associated disturbances such as anxiety, were corrected, and brain Aβ levels significantly ameliorated. AIK3a305 may therefore be a promising novel therapeutic strategy for AD patients.

## Introduction

Despite decades of intense research and recent regulatory approvals of new biological-based treatments (Walsh et al., 2024), disease-modifying therapeutic approaches for Alzheimer’s disease (AD) are still very much needed (Sarazin et al., 2024), since current treatments are not satisfactory (Richard et al., 2021; Atwood and Perry, 2023). Moreover, new therapies are centered in treating AD at very early stages, as repeated failure of clinical trials in even modestly advanced AD cases has led to the prevailing notion that when clinical symptoms debut, the pathological process is so advanced that it cannot longer be reverted (Rasmussen and Langerman, 2019). For this reason, even moderate AD can be considered a neglected disease, although millions worldwide are affected (2021).

IGF-1 is a pleiotropic neurotrophic factor with wide neuroprotective activity (Fernandez and Torres-Aleman, 2012), Altered IGF-1 levels have been reported in the brain in the majority of neurological disorders (Trejo et al., 2004), suggesting that dysregulation of IGF-1 signaling is a common feature of brain pathology and may contribute to disease progression. In AD, we postulated that loss of IGF-1 function due to reduced IGF-1 receptor sensitivity, IGF-1 deficiency, or both, favors development of AD-related pathogenic events (Carro and Torres-Aleman, 2004). Since inflammation, excitotoxicity, and oxidative stress associated to AD (Behl et al., 1994; Akiyama et al., 2000; Ittner et al., 2010) elicit IGF-1 resistance (Venters et al., 1999; Garcia-Galloway et al., 2003; Davila and Torres-Aleman, 2008), and a variety of observations repeatedly indicate that IGF-1 participates in AD pathology (Moloney et al., 2010; de Bruijn et al., 2014; Zegarra-Valdivia et al., 2023; Zhang et al., 2026), in the present study we used AIK3a305, a recently developed IGF-1 sensitizer (Zegarra-Valdivia et al., 2022b), to determine its potential therapeutic utility in this condition. Significantly, 1-year-old APP/PS1 mice, a model of familial AD that at this age already develop clear symptoms of the disease (Borchelt et al., 1997), show normal cognition and mood, and reduced brain amyloid β (Aβ) levels, after 3 months of daily treatment with AIK3a305.

## Materials and Methods

### Animals

Wild type (C57BL/6JolaHsd; 12 months old) and APP_swe_ and PS1Δ9 mice of C57BL6/J background (APP/PS1; 12 months-old), a kind gift of P. Mouton (NIH), were used. Mice were maintained according to ARRIVE guidelines as indicated before (Zegarra-Valdivia et al., 2022a). Animals were housed in standard cages (48 × 26 cm^2^) with 5 per cage, and kept on a light-dark cycle (12-12 h, lights on at 8 am) at constant temperature (22°C) and humidity, and with food (pellet rodent diet) and water *ad libitum*. All experimental protocols were performed during the light cycle and followed European guidelines (86/609/EEC & 2003/65/EC, European Council Directives). Studies were approved by the local Bioethics Committee (UPV M20). Animals were not randomized and were used in a sex-balanced manner throughout. Potential confounders were not accounted for. Each experimenter took account of group allocation under study. All efforts were done to reduce harm to the animals. Mice were handled for three days prior to any experimental manipulations and familiarized with behavioral arenas to minimize novelty stress, or deeply anesthesized with pentobarbital prior to sacrifice, when needed. Sample sizes were kept as little as possible to comply with current animal reduction policies, using at least 10 animals per group, unless otherwise indicated. No adverse events were expected, nor found. End-point measures included checking reflexes in deeply anesthesized animals prior to culling.

### Materials

We used a novel IGF-1 sensitizer, AIK3a305 (Allinky Biopharma, Spain), as described before (Zegarra-Valdivia et al., 2022b). AIK3a305 or the vehicle (DMSO + saline) were injected intra-peritoneally (ip) daily for 3 months in 1-year-old APP/PS1 mice. This compound shows good blood-brain-barrier penetration and enhances responses to IGF-1 (Zegarra-Valdivia et al., 2022b). We controlled gain in body weight along the 3 months of the study and found no effects of AIK3a305 administration. This compound has already been shown to have no toxicity in rats and dogs (unpublished observations).

### Behavioral tests

#### Elevated plus maze

To assess anxiety-like/coping behavior, mice were introduced in a maze of 40 cm from the floor with two opposing arms. Two protected (closed) arms (30 cm (length) × 5 cm (wide) × 15.25 (height), and two opposing unprotected (open) arms (30 cm (length) × 5 cm (wide). Each animal was introduced in the middle of the apparatus for 5 minutes. Stress was scored as time spent in the closed arms while coping behavior was estimated by time spent in the open arms. All measures were recorded (Video Tracking Plus Maze Mouse; Med Associates, USA), and analyzed as described (Munive et al., 2019).

#### Y maze

This test measures spontaneous alternation as an index of working memory (Sarter et al., 1988) and was used as described (Zegarra-Valdivia et al., 2022a). In brief, the mouse is placed at the end of one arm to move freely from side to side of the maze during an 8-min session. Videos recorded the sequence of entries during the whole time of the experiment and were analyzed off-line. Entrance to each arm is scored when the mouse places the hind paws entirely in the zone. Alternation was defined as successive entries into the three arms on overlapping triplet sets. Consecutive triplets were analyzed, and alternate behavior was calculated as the percentage of actual alternation (number of triplets with non-repeated entries) versus total alternation opportunities (total number of triplets), as described (Recinto *et al*., 2012; Yan *et al*., 2017).

### ELISA

Aβ_1-40_ levels were determined in brain lysates using commercial immunoassays (ThermoFisher. Ref # KHB3481) following the manufacturer’s instructions.

### Statistics

Statistical analysis was performed using GraphPad Prism 6 software (San Diego, CA, USA). Depending on the number of independent variables, normally distributed data (Kolmogorov-Smirnov normality test), and the experimental groups compared, we used either two-way ANOVAs, or Two-way repeated measure ANOVA, followed by Sidak’s multiple comparison test. For non-normally distributed data, we used the Mann-Whitney U test to compare two groups, Kruskal-Wallis as a Post Hoc analysis. The sample size for each experiment was chosen based on previous experience and aimed to detect at least a p<0.05 in the different tests applied, considering a reduced use of animals. Results are shown as mean ± standard error (SEM) and *p* values coded as follows: *p< 0.05, **p< 0.01, ***p< 0.001. Animals were included in each experimental group randomly by the researcher. No animals were excluded from the analyses.

## Results

One-year old APP/PS1 mice were treated daily for 3 months with AIK3a305 (Figure 1A) and behavior was tested at 1, and 3 months during treatment (Figure 1A). As early as 1 month later, working memory, assessed in the Y maze, was normalized by AIK3a305 treatment, and kept normal for the duration of the study (Figure 1B). Anxiety, measured in the elevated plus maze, was normalized in APP/PS1 mice after 3 months of treatment with AIK3a305 (Figure 1C). However, social behavior alterations were not normalized by treatment, as seen in the social novelty and social affiliation tests (not shown). Finally, elevated levels of Aβ_1-40_ were significantly diminished by AIK3a305 treatment (Figure 1D).

**Figure 1:**
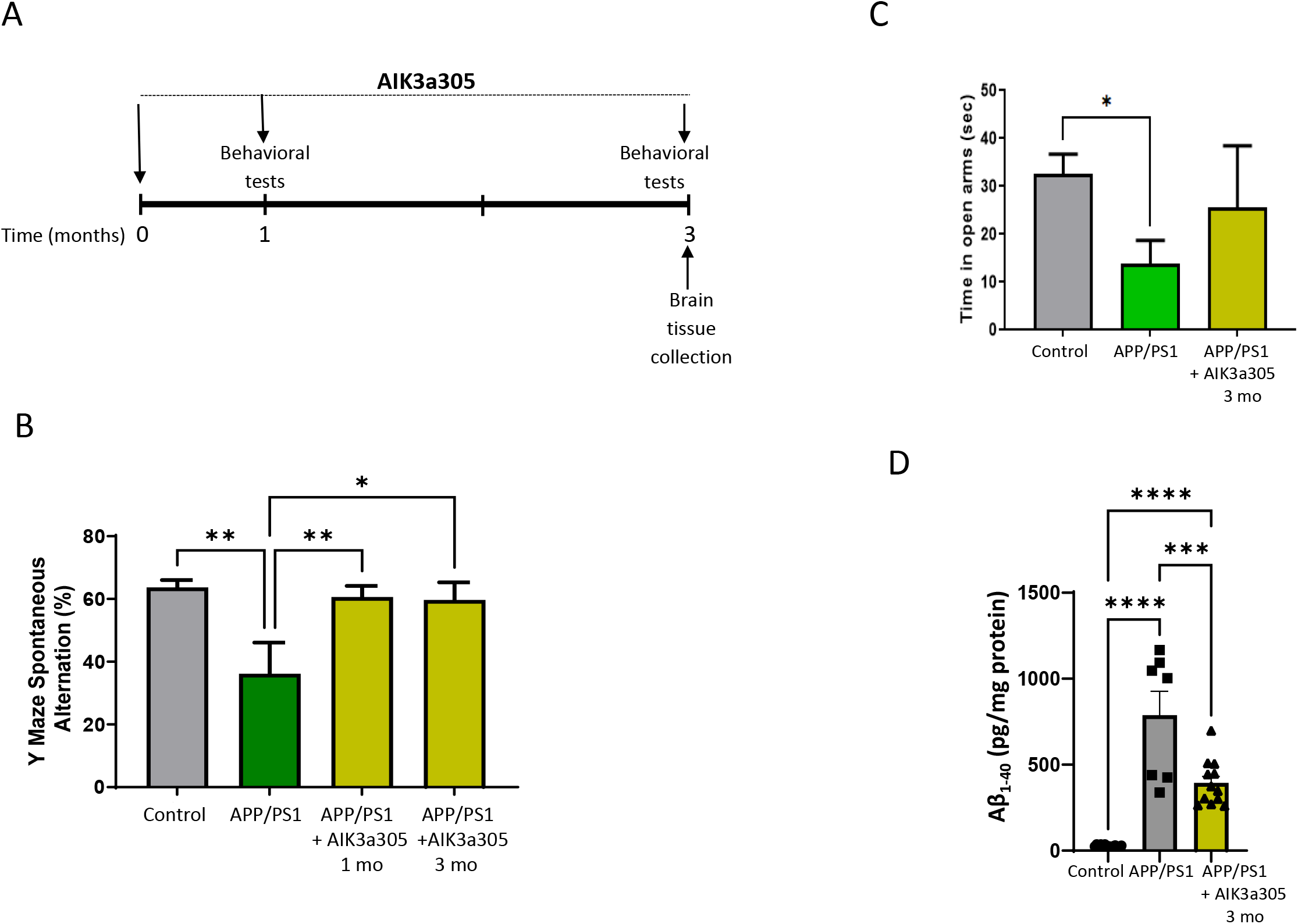
AD and AIK3a305. **A**, Time-line of treatment with AIK3a305 in 1-year old APP/PS1 mice. **B**, Treatment with AIK3a305 normalized working memory in the Y maze 1 month after starting treatment, and lasted for the duration of the study (until 3 months). **C**, AIK3a305 normalized anxiety levels, as seen in the elevated plus maze at 3 months. **D**, Treatment with AIK3a305 significantly reduced brain Aβ_1-40_ levels after 3 months.

## Discussion

These results indicate that treatment of 1-year-old APP/PS1 mice with AIK3a305, an IGF-1 sensitizer, that also displays various others neuroactive (Zegarra-Valdivia et al., 2022b; Zegarra-Valdivia et al., 2026), and therapeutic activities (unpublished observations), ameliorates major neurological disturbances, including impaired cognition and higher anxiety, and significantly decreases brain Aβ_1-40_ levels, supporting a potential therapeutic role of IGF-1 in this pathology.

From the clinical point of view, the most severe effect of AD is loss of cognitive abilities such as working memory, that substantiates daily life activities (Kirova et al., 2015). In APP/PS1 mice, AIK3a305 treatment normalized it, together with anxiety, a mood alteration commonly associated to dementia (Botto et al., 2022). Intriguingly, social alterations usually associated to AD-like amyloidosis in APP/PS1 mice were not corrected by the IGF-1 sensitizer. Reinforcing the potential role of AIK3a305 as a novel therapeutic tool in AD, elevated brain levels of Aβ, a pathological hallmark of the disease were also significantly decreased.

The potential utility of IGF-1 for treatment of AD has been well documented (Carro et al., 2002; Dunacka et al., 2025), although apparently contradictory data questioned whether it may be useful for this and other neurological conditions since in many reports (Cohen et al., 2009; Freude et al., 2009; El-Ami et al., 2014; Gontier et al., 2015), but not in all (Narayan et al., 2025), partial genetic ablation of the IGF-1 receptor showed to be neuroprotective. As in our hands, but not in others’ (Lanz et al., 2008), IGF-1 has been always neuroprotective after its administration in experimental models (Fernandez et al., 1998; Carro et al., 2006) and human patients (Arpa et al., 2011), we developed a small molecule IGF-1 sensitizer able to cross the BBB, to circumvent regulatory issues. The recent publication that a small IGF-1 mimetic (Trofinetide), recently approved for treatment of several autistic spectrum disorders (Neul et al., 2023), and shown efficacious in diverse neurological diseases (Grunseich et al., 2018), proved to be effective in APP/PS1 mice (Chen et al., 2025), further supports that potentiation of IGF-1 activity in AD-like, and other brain pathologies is a promising therapeutic approach. Since patients with Alzheimer’s disease currently have no effective treatment options for cognitive decline, testing AIK3a305 in this population should be considered—provided that the compound receives approval for human use.

Phase 1 clinical trials are already in progress.

## Acknowledgements

We thank the excellent work of Achucarro Core Facilities, and in particular of R Cipriani.

## References

(2021) 2021 Alzheimer’s disease facts and figures. Alzheimer’s & Dementia 17:327–406.

Akiyama H et al. (2000) Inflammation and Alzheimer’s disease. Neurobiol Aging 21:383–421.

Arpa J, Sanz-Gallego I, Medina-Baez J, Portela LV, Jardim LB, Torres-Aleman I, Saute JA (2011) Subcutaneous insulin-like growth factor-1 treatment in spinocerebellar ataxias: an open label clinical trial. Mov Disord 26:358–359.

Atwood CS, Perry G (2023) Playing Russian Roulette with Alzheimer’s Disease Patients: Do the Cognitive Benefits of Lecanemab Outweigh the Risk of Edema, Stroke and Encephalitis? Journal of Alzheimer’s Disease 92:799–801.

Behl C, Davis JB, Lesley R, Schubert D (1994) Hydrogen peroxide mediates amyloid beta protein toxicity. Cell 77:817–827.

Borchelt DR, Ratovitski T, van Lare J, Lee MK, Gonzales V, Jenkins NA, Copeland NG, Price DL, Sisodia SS (1997) Accelerated amyloid deposition in the brains of transgenic mice coexpressing mutant presenilin 1 and amyloid precursor proteins. Neuron 19:939–945.

Botto R, Callai N, Cermelli A, Causarano L, Rainero I (2022) Anxiety and depression in Alzheimer’s disease: a systematic review of pathogenetic mechanisms and relation to cognitive decline. Neurol Sci 43:4107–4124.

Carro E, Torres-Aleman I (2004) The role of insulin and insulin-like growth factor I in the molecular and cellular mechanisms underlying the pathology of Alzheimer’s disease. Eur J Pharmacol 490:127–133.

Carro E, Trejo JL, Gomez-Isla T, LeRoith D, Torres-Aleman I (2002) Serum insulin-like growth factor I regulates brain amyloid-beta levels. Nat Med 8:1390–1397.

Carro E, Trejo JL, Gerber A, Loetscher H, Torrado J, Metzger F, Torres-Aleman I (2006) Therapeutic actions of insulin-like growth factor I on APP/PS2 mice with severe brain amyloidosis. Neurobiol Aging 27:1250–1257.

Chen M, Ning Y, Yang H, Jia J (2025) Trofinetide Improves Cognitive Function in APP/PS1 Mice by Suppressing Inflammation and Apoptosis. Mol Neurobiol 63:129.

Cohen E, Paulsson JF, Blinder P, Burstyn-Cohen T, Du D, Estepa G, Adame A, Pham HM, Holzenberger M, Kelly JW, Masliah E, Dillin A (2009) Reduced IGF-1 signaling delays age-associated proteotoxicity in mice. Cell 139:1157–1169.

Davila D, Torres-Aleman I (2008) Neuronal Death by Oxidative Stress Involves Activation of FOXO3 through a Two-Arm Pathway That Activates Stress Kinases and Attenuates Insulin-like Growth Factor I Signaling. Molecular Biology of the Cell 19:2014–2025.

de Bruijn RF, Janssen JA, Brugts MP, van Duijn CM, Hofman A, Koudstaal PJ, Ikram MA (2014) Insulin-Like Growth Factor-I Receptor Stimulating Activity is Associated with Dementia. J Alzheimers Dis.

Dunacka J, Grembecka B, Majkutewicz I, Wrona D (2025) Central Insulin-like Growth Factor-1 Treatment Enhances Working and Reference Memory by Reducing Neuroinflammation and Amyloid Beta Deposition in a Rat Model of Sporadic Alzheimer’s Disease. Pharmaceuticals (Basel) 18.

El-Ami T, Moll L, Carvalhal Marques F, Volovik Y, Reuveni H, Cohen E (2014) A novel inhibitor of the insulin/IGF signaling pathway protects from age-onset, neurodegeneration-linked proteotoxicity. Aging Cell 13:165–174.

Fernandez AM, Torres-Aleman I (2012) The many faces of insulin-like peptide signalling in the brain. Nat Rev Neurosci 13:225–239.

Fernandez AM, de la Vega AG, Torres-Aleman I (1998) Insulin-like growth factor I restores motor coordination in a rat model of cerebellar ataxia. Proc Natl Acad Sci U S A 95:1253–1258.

Freude S, Hettich MM, Schumann C, Stohr O, Koch L, Kohler C, Udelhoven M, Leeser U, Muller M, Kubota N, Kadowaki T, Krone W, Schroder H, Bruning JC, Schubert M (2009) Neuronal IGF-1 resistance reduces A{beta} accumulation and protects against premature death in a model of Alzheimer’s disease. FASEB J 23:3315–3324.

Garcia-Galloway E, Arango C, Pons S, Torres-Aleman I (2003) Glutamate excitotoxicity attenuates insulin-like growth factor-i prosurvival signaling. Mol Cell Neurosci 24:1027–1037.

Gontier G, George C, Chaker Z, Holzenberger M, Aid S (2015) Blocking IGF Signaling in Adult Neurons Alleviates Alzheimer’s Disease Pathology through Amyloid-beta Clearance. J Neurosci 35:11500–11513.

Grunseich C, Miller R, Swan T, Glass DJ, El Mouelhi M, Fornaro M, Petricoul O, Vostiar I, Roubenoff R, Meriggioli MN, Kokkinis A, Guber RD, Budron MS, Vissing J, Soraru G, Mozaffar T, Ludolph A, Kissel JT, Fischbeck KH, group BVSs (2018) Safety, tolerability, and preliminary efficacy of an IGF-1 mimetic in patients with spinal and bulbar muscular atrophy: a randomised, placebo-controlled trial. Lancet Neurol 17:1043–1052.

Ittner LM, Ke YD, Delerue F, Bi M, Gladbach A, van EJ, Wolfing H, Chieng BC, Christie MJ, Napier IA, Eckert A, Staufenbiel M, Hardeman E, Gotz J (2010) Dendritic function of tau mediates amyloid-beta toxicity in Alzheimer’s disease mouse models. Cell 142:387–397.

Kirova AM, Bays RB, Lagalwar S (2015) Working memory and executive function decline across normal aging, mild cognitive impairment, and Alzheimer’s disease. Biomed Res Int 2015:748212.

Lanz TA, Salatto CT, Semproni AR, Marconi M, Brown TM, Richter KE, Schmidt K, Nelson FR, Schachter JB (2008) Peripheral elevation of IGF-1 fails to alter Abeta clearance in multiple in vivo models. Biochem Pharmacol 75:1093–1103.

Moloney AM, Griffin RJ, Timmons S, O’Connor R, Ravid R, O’Neill C (2010) Defects in IGF-1 receptor, insulin receptor and IRS-1/2 in Alzheimer’s disease indicate possible resistance to IGF-1 and insulin signalling. Neurobiology of Aging 31:224–243.

Munive V, Zegarra-Valdivia JA, Herrero-Labrador R, Fernandez AM, Aleman IT (2019) Loss of the interaction between estradiol and insulin-like growth factor I in brain endothelial cells associates to changes in mood homeostasis during peri-menopause in mice. Aging 11:174–184.

Narayan S, Mao K, Williams-Medina AR, Richmann T, Gal M, Engel M, Zhang Y, Graff S, Sidoli S, Barzilai N, Huffman DM (2025) Reduced IGF-1 signaling fails to limit Alzheimer’s disease progression in a novel rat model of IGF-1R haploinsufficiency. Sci Rep 16:1856.

Neul JL, Percy AK, Benke TA, Berry-Kravis EM, Glaze DG, Marsh ED, Lin T, Stankovic S, Bishop KM, Youakim JM (2023) Trofinetide for the treatment of Rett syndrome: a randomized phase 3 study. Nat Med.

Rasmussen J, Langerman H (2019) Alzheimer’s Disease - Why We Need Early Diagnosis. Degener Neurol Neuromuscul Dis 9:123–130.

Richard E, den Brok M, van Gool WA (2021) Bayes analysis supports null hypothesis of anti-amyloid beta therapy in Alzheimer’s disease. Alzheimers Dement 17:1051–1055.

Sarazin M, Lagarde J, El Haddad I, de Souza LC, Bellier B, Potier M-C, Bottlaender M, Dorothée G (2024) The path to next-generation disease-modifying immunomodulatory combination therapies in Alzheimer’s disease. Nature Aging 4:761–770.

Sarter M, Bodewitz G, Stephens DN (1988) Attenuation of scopolamine-induced impairment of spontaneous alteration behaviour by antagonist but not inverse agonist and agonist beta-carbolines. Psychopharmacology (Berl) 94:491–495.

Trejo JL, Carro E, Garcia-Galloway E, Torres-Aleman I (2004) Role of insulin-like growth factor I signaling in neurodegenerative diseases. J Mol Med 82:156–162.

Venters HD, Tang Q, Liu Q, VanHoy RW, Dantzer R, Kelley KW (1999) A new mechanism of neurodegeneration: a proinflammatory cytokine inhibits receptor signaling by a survival peptide. Proc Natl Acad Sci U S A 96:9879–9884.

Walsh S, Merrick R, Milne R, Nurock S, Richard E, Brayne C (2024) Considering challenges for the new Alzheimer’s drugs: Clinical, population, and health system perspectives. Alzheimers Dement.

Zegarra-Valdivia J, Fernandez AM, Martinez-Rachadell L, Herrero-Labrador R, Fernandes J, Torres Aleman I (2022a) Insulin and insulin-like growth factor-I receptors in astrocytes exert different effects on behavior and Alzheimer s-like pathology. F1000Res 11:663.

Zegarra-Valdivia JA, Pignatelli J, Nuñez A, Torres Aleman I (2023) The Role of Insulin-like Growth Factor I in Mechanisms of Resilience and Vulnerability to Sporadic Alzheimer’s Disease. Int J Mol Sci 24.

Zegarra-Valdivia JA, Khan MZ, Putzolu A, Pignatelli J, Torres Aleman I (2026) Therapeutic Effects of An Insulin-Like Growth Factor I Sensitizer In Traumatic Brain Injury. bioRxiv:2026.2005.2013.724506.

Zegarra-Valdivia JA, Fernandes J, Fernandez de Sevilla ME, Trueba-Saiz A, Pignatelli J, Suda K, Martinez-Rachadell L, Fernandez AM, Esparza J, Vega M, Nunez A, Aleman IT (2022b) Insulin-like growth factor I sensitization rejuvenates sleep patterns in old mice. Geroscience 44:2243–2257.

Zhang J, He Y, Yang P, Zhang H, Tong Y, Jiang L, Li Z, Yan M, Li X, Yang Q, Yang J, Yuan Z, Zhang J, Cheng J (2026) Microglial Rack1 Deficiency Alleviates Alzheimer’s Disease Pathology through Enhancing IGF1-Mediated Astrocytic Phagocytosis. Adv Sci (Weinh) 13:e15877.

